# Cell-type-agnostic differential gene expression uncovers conserved principles of cellular regulation

**DOI:** 10.1101/2025.09.29.679285

**Authors:** Alexander R Hummels, Carlos Camacho

## Abstract

Cellular responses to perturbagens—including pharmacological compounds and genetic manipulations—remain incompletely characterized, as conventional approaches are constrained by cell-type-specific biases. Here, we derive consensus signatures (CS) that uncover conserved principles not of gene expression, but of fundamental cellular regulation. Quantified by aggregating differential gene expression responses across diverse cell types, time points, and doses, these CS retain the core features of individual experiments while preserving biological relevance. As proof of concept, genome-wide CS screening identified three knockdown-resistant genes—CCNA1, ORC1, and SOX2—that also engage in their reciprocal up-regulation. Expression of this network is modulated by the factor E2F2, revealing a mechanism by which proliferative signaling persists despite diverse genetic and chemical perturbations. Crucially, we identify that ORC1 down-regulation by specific CDK4/6 inhibitors singularly disrupts this feedback loop, providing a direct route to suppress E2F2-driven proliferation. More generally, we demonstrate for the first time that CS can accurately predict RNA-seq responses in novel cell lines, uncovering evolutionarily conserved mechanisms that regulate fundamental biological processes beyond context-specific variability.

---

Cellular responses to perturbations are governed by evolutionarily conserved regulatory mechanisms. While perturbation biology has long relied on data with shared cellular contexts – from early developments in biclustering applied to yeast microarrays (1–5) to more recent tissue-specific coexpression network analyses (6) – this approach inherently limits generalizability across cell types. Although context-specific analyses yield valuable insights, they rarely extrapolate to other biological systems.

There is growing evidence that integrating data from different contexts has predictive power. Bulk cell RNA-seq, which implicitly integrates the responses of multiple cells with different spatial and environmental factors, has been successfully combined with single-cell RNA-seq (scRNA-seq) in predicting cancer drug response, reconstructing cell-line-specific gene regulatory networks, and imputing genes subject to dropout events (7–10). scRNA-seq data has also been explicitly averaged in post-processing, combining cell-specific indicators to provide more robust and predictive estimation of global co-expression (11). These studies demonstrate the power of moving from highly specific contexts to broader relevance, but only operate in a cell-type-specific niche and do not generalize to other cell types. More recently, analysis of publicly available tissue samples has revealed conserved transcriptional modules between both normal and cancer tissues of various types (12). This striking transcriptional similarity indicates that context integration may indeed be possible at the level of tissues or cell lines.

Emerging evidence indicates that perturbation biology approaches can facilitate data integration across diverse cellular contexts, enabling the derivation of generalizable biological insights. One study found 70% of drug-induced functional modules were conserved across three human cancer cell lines (13). Another averaged drug response profiles over five cell lines to reveal nearly 50 cell-line-independent functional modules (14). More recently, the LINCS L1000 dataset (15) - encompassing mRNA signatures from nearly 40,000 small-molecule treatments and over 4,000 gene knockdowns across more than 100 cell lines - has enabled countless more opportunities for data integration and downstream inference (16–22). One example, Pabon et al., utilized Z-scores from four or more cell lines to create a robust transcriptional response model that facilitates drug-target interaction identification (18). Collectively, these findings strongly suggest that cross-context integration of large-scale datasets can significantly enhance our understanding of fundamental cellular processes.

In this study, we first establish that perturbagen-induced differential expression exhibits remarkable consistency across cellular contexts, enabling the construction of robust consensus signatures. We then provide experimental validation that these consensus signatures accurately predict treatment responses in novel cellular environments. Leveraging the genome-wide consensus signatures obtained from LINCS dataset, we show that knockdown-resistant genes are functionally enriched in cell-cycle regulation, revealing how proliferative signaling persists despite diverse genetic and chemical perturbations. Equally significant, our findings reveal ORC1 (origin of replication subunit 1) as a critical mediator of both positive and negative feedback loops controlling E2F regulation, a previously unrecognized mechanism exploited by clinically approved inhibitors. By employing a hypothesis-free framework, perturbation biology enables robust and generalizable analysis of biological processes across diverse cell-types.

## Results

### Perturbagen-induced differential gene expression shows consistency across cell types

Given previous work demonstrating the conservation of drug-induced functional modules across cell types, we investigated whether perturbagens response profiles could be validated directly as context-independent. To ensure analytical rigor, we adhered to LINCS’ gold-standard requirements for reproducible and distinct signatures. For all perturbagens with gold-standard signatures in at least 2 of the 9 LINCS core cell lines, we measured perturbagen-specific cross-cell consistency controlling for time, dose, and shRNA sequence (where applicable). Notably, LINCS’ gold-standard consistency criterion is that the 75th percentile of pairwise replicate Spearman correlations should be at least 0.2, ensuring that replicates represent reproducible signatures. We extend this test to cross-cell comparisons for experiments with the same perturbagen, time, and dose but different cells to investigate whether perturbagens elicit reproducible responses across cellular contexts.

For knockdowns (KDs), we find a majority of shRNA perturbations exhibit gold-standard consistency across cell lines at all data levels (Fig. 1A). However, Level 4 and 5 data for shRNAs correspond to individual shRNA sequences, which are well-known to have strong seed-mediated off-target effects in general (23–25). Further, these off-target effects have been demonstrated to be more well conserved than their on-target effects in the LINCS dataset (15). While these considerations complicate the interpretation of the consistency demonstrated at Levels 4 and 5, Level 6 consensus genetic signatures (CGS) LINCS’ primary method for deriving on-target shRNA profiles by averaging over shRNA sequences demonstrates remarkable on-target consistency across cell lines. After filtering off-target effects from these signatures, knockdowns soundly demonstrate that perturbation responses can be conserved across contexts.

**Fig. 1.**
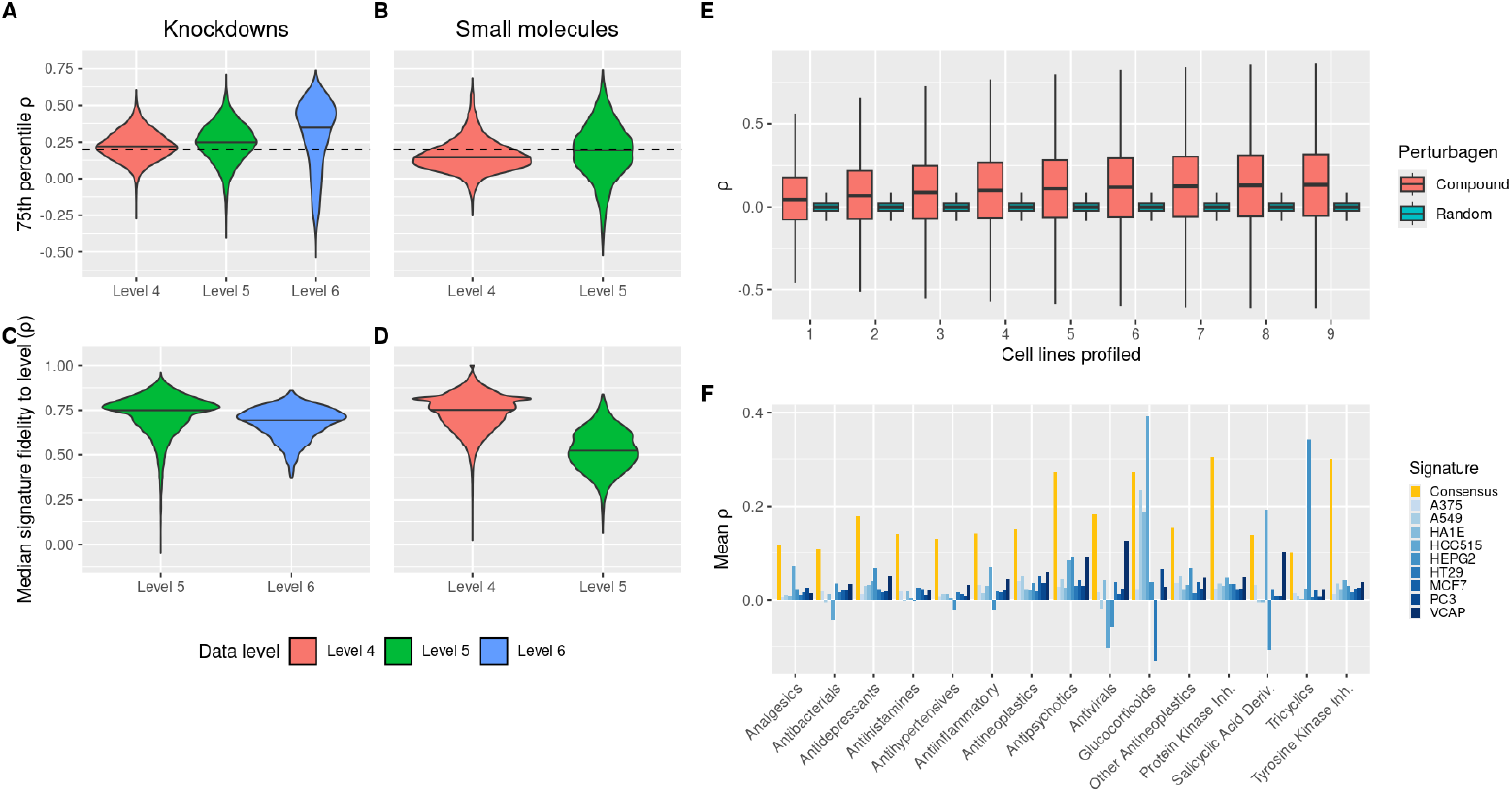
Consensus perturbagen signatures (CS) reveal consistency across cell lines. A) 75th percentile of pairwise shRNA KD inter-cell correlation of Level 4 (replicate), Level 5 (signature), and Level 6 (consensus genetic signature [CGS]) data for KDs profiled in at least 2 cell lines, controlling for time, dose, and sequence (Levels 4, 5). B) 75th percentile of pairwise small molecule inter-cell correlation of Level 4 (replicate) and Level 5 (signature) data for small molecules profiled in at least 2 cell lines, controlling for time and dose. C) shRNA CGS signature median correlation to component Level 5 signatures (left; median 3 signatures per CGS) and KD CS median correlation to component Level 6 CGS signatures (right; median 5 CGS signatures per CS). D) Small molecule signature median correlation to component Level 4 replicates (left; median 3 replicates per signature) and CS median correlation to component Level 5 signatures (right; median 10 signatures per CS). E) Inter-perturbagen correlation for CS and random signatures utilizing varying numbers of cell lines. F) Mean intra-class correlation of annotated drugs by single cell line or all-cell consensus. All correlations Spearman over LM genes. Dashed line in A, B is gold-standard consistency threshold criterion from LINCS (75th percentile pairwise Spearman correlation *≥* 0.2).

Small molecules by comparison show less cross-cell consistency across data levels. Only 30% of small molecules show gold-standard consistency at the replicate level, but this improves to nearly 50% at the signature level, amounting to over 3,000 small molecules (Fig. 1B). Part of this lower consistency is explained by the known lower transcriptional activity of small molecules when compared to knockdowns (revisited later) (15). Importantly, the small molecules considered here include thousands of drug-like molecules from the BROAD institute - small molecules with no confirmed targets-that may not have strong affinity for any target. The above notwithstanding, this increase in signal at the signature level validates the ability of moderated Z-scores (MODZ, LINCS’ method for creating signatures from a collection of experimental profiles) to accurately capture the underlying biology of a group of experiments.

Systematic evaluation further demonstrates even better signature similarity across doses and observation time-scales (Fig. S3A-D), suggesting that experimental signatures can be integrated over a wide variety of experimental factors. In total, over 4,000 small molecule and shRNA knockdowns demonstrate transcriptional profiles that maintain sufficient consistency to support the creation of consensus signatures through integration across diverse cellular contexts, dosage regimes, and temporal conditions. We emphasize that previous studies that averaged drug response across cell lines (14) do not provide this important cross-check that signatures they are compiling represent a reproducible cellular signature. Further, our results suggest that a large proportion (50%) of drugs do not elicit a consistent cellular response, complicating downstream analyses of averaged responses without this validation.

### Consensus signatures are cell-type agnostic

We derived consensus perturbagen signatures (CS) following the analytical pipeline detailed in Methods, adopting the goldstandard quality control metrics employed by the LINCS consortium. To verify that CS are representative of profiled contexts, we computed the median Spearman correlation between each CS and its different (cell lines, doses, times) constituent experimental signatures. Remarkably, this similarity for KD CS is distinctly similar to that of LINCS single gene CGS (Fig. 1C), despite the tougher challenge of incorporating greater heterogeneity in cells, times and doses (median 5 signatures for KD CS, 3 for CGS). We also observe that despite the lower transcriptional activity of small molecules, level 4 replicates can be compiled into a single representative, reproducible signatures much like shRNA knockdowns (Fig. 1D). While small-molecule CS are not as strongly representative, a median Spearman correlation of 0.5 is substantial given an even higher degree of heterogeneity on top of previous concerns about transcriptional activity (median 10 signatures for CS, 3 replicates for a regular signature).

### Integrating four or more cell lines amplifies signal in consensus signatures

As our stated goal is to interrogate cell-type-agnostic response, it is crucial to ask: at what point do the CS above no longer represent any individual cell line but instead an agnostic regulatory response? We have hypothesized that specific contexts most notably cell lines - blur notions of general inter-perturbagen similarity or dissimilarity (see Methods, Eq. (2) and Eq. (3)). Thus, we measured inter-perturbagen similarity for CS created from varying numbers of cell lines to see at what point adding cell lines no longer amplified these signals. Particularly, we considered 557 small molecules profiled in all 9 core LINCS cell lines. shRNA knockdowns are not considered, as only 5 knockdowns have gold-standard CGS signatures in all 9 cell lines. For each compound and all 512 cell line combinations, we computed unique CS and evaluated inter-compound relationships using Spearman correlation as before (Fig. 1E). As control, we verified that MODZ did not homogenize random signatures devoid of biological content (see Methods).

This analysis reveals that the inter-compound similarity distribution broadens as more cell lines are incorporated into the CS. This trend is conserved when integrating increasing doses or timepoints instead of cell lines (Fig. S3E-F, see **??**). While there is no clear point of diminishing returns, Fig. 1E suggests 4 cell lines as the inflection point for distribution mean stabilization. This threshold aligns with prior findings that 4 cell lines suffice to detect meaningful correlational patterns between genetic and pharmacological perturbations (18). Consequently, we restrict all subsequent analyses to CS derived from at least 4 cell lines.

### Consensus signatures enhance similarity detection within pharmacological classes

Since CS systematically enhance general inter-perturbagen similarity, we hypothesized they should particularly amplify similarities among pharmacologically related compounds. We tested this hypothesis using 15 well-annotated drug classes from the LINCS data portal (26), comparing intra-class similarity between single-cell signatures and multi-cell CS. Remarkably, CS showed stronger intra-class similarity than any single cell line in 12 of 15 classes (Fig. 1H). Formal statistical analysis using one-sided Mann-Whitney U tests confirmed that CS capture significantly more intra-class similarity than any individual cell line for 8 of 15 classes. Across all comparisons, CS outperformed single-cell signatures in 85% of cases (Table S1), demonstrating their superior ability to resolve pharmacological relationships.

### Transcriptional activity dictates consensus signature reliability

While 2,663 small molecule CS and 1,002 gene KD CS meet LINCS’ gold-standard thresholds, 40% of perturbagens profiled in at least four cell lines do not. We hypothesized that these perturbagens exhibited reduced activity, as lower signal-to-noise ratios would prevent consistent detection across individual experiments, precluding a reliable consensus signature derivation. Indeed, comparing the transcriptional activity score (TAS) of experiments for perturbagens with gold-standard and non-gold-standard CS, we find a highly significant difference. Specifically, nearly 90% of CS that do not reach LINCS gold-standard consistency have more weak signatures (TAS ≤ 0.2) than strong signatures (Fig. 2A), and over 96% of perturbagens that don’t reach gold-standard consensus are small molecules.

**Fig. 2.**
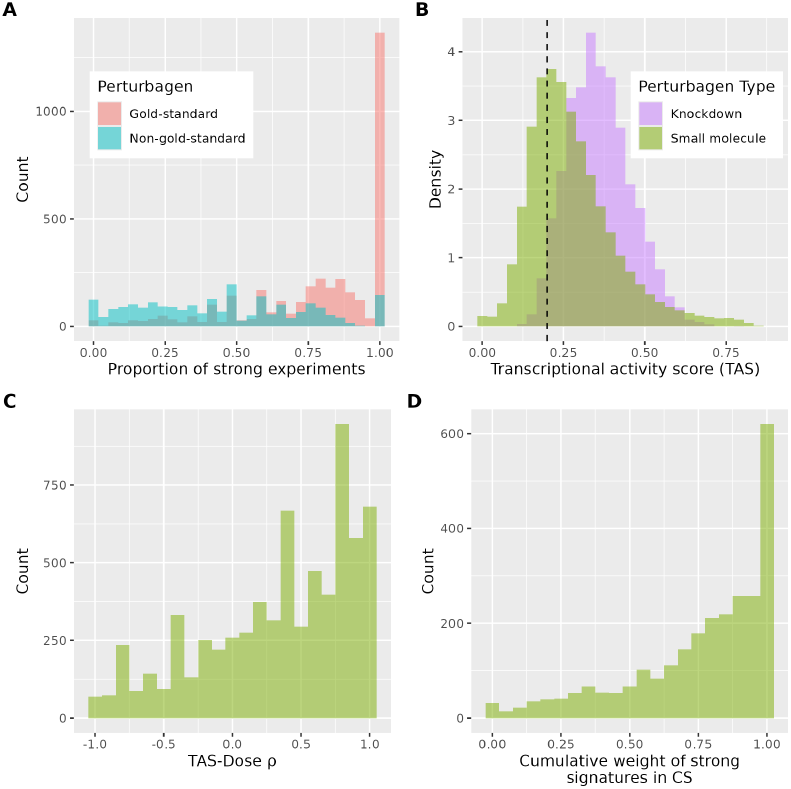
Transcriptional activity dictates CS reliability. A) Proportion of individual experiments considered strong (TAS > 0.2) for gold-standard, non-gold-standard perturbagen CS. Gold, non-gold perturbagens vary significantly on this proportion (Mann-Whitney *p <* 2.23 *×* 10^*−*308^, smallest floating point number for R). B) Individual experiment TAS of gold-standard small molecules and knockdowns. High activity threshold (0.2) represented with dashed line. C) TAS and dosage positively correlate for small molecules with at least 4 doses when controlling for cell line, timepoint. D) Cumulative weight of strong signatures (TAS > 0.2) in small-molecule CS. Most CS predominantly composed of high-activity signatures with high weighting from MODZ.

Small molecules have been reported to have a lower transcriptional activity than shRNAs (15), and we recapitulate this finding for perturbagens with gold-standard CS (Fig. 2B). This explains why a smaller proportion of small molecules achieve gold-standard consistency across cell lines when compared to knockdowns in Fig. 1A-B. Additionally, nearly half of signatures comprising small-molecule CS have weak transcriptional activity, further explaining why small molecule CS have decreased similarity to their component signatures (Fig. 1D). Indeed, small molecules induce variable, context-dependent signaling that inherently reduces signal-to-noise ratios, whereas shRNA KD produce binary effects by eliminating target proteins. This distinction reflects the reality that signaling networks are disrupted by small molecules and circumvented by KD, making representative pharmacological CS more challenging to derive than genetic CS.

Unsurprisingly, we find that this notion of transcriptional activity has a generally positive correlation with dose, suggesting that at low enough doses the cellular response of some compounds may get reduced to noise (Fig. 2C). This clarifies our earlier observations about cell-type-agnostic response, as perturbagens must be present in a high enough concentration to force the cell to react. But if this condition is met, we see that thousands of small molecule and shRNA KD perturbagens activate the same response across cell lines.

Finally, while we do not exclude weak signatures from our CS, we hypothesized that these signatures would receive little weight from MODZ and thus be drowned out by strong signatures. Indeed, we find the cumulative weight of these weak experiments across out 2,663 small-molecule CS to be generally insignificant (Fig. 2D). This combined with the observation that effect size is explicitly part of the TAS computation, means that low-TAS signatures have lower absolute Z scores and are given small weight. This suggests that weak signatures do not appreciable bias our CS, and that CS represent cell-type agnostic cellular responses to perturbagens at an effective dose.

### Consensus signatures enable cross-cell-line prediction of drug responses in unprofiled cellular contexts

One of the most powerful applications of a genome-wide CS is its capacity to serve as a comprehensive reference database, enabling rapid prediction of responses across hundreds of genes and thousands of related perturbations. To validate the predictive capabilities of our CS for unprofiled cellular contexts, we analyzed the CS of STL (a FOXM1 degrader, (20)) against independent RNA-seq data from Raghuwanshi et al. (21) profiling STL001, a chemical analog of STL, in two cell lines: FLO1 (esophageal adenocarcinoma) and OVCAR8 (ovarian carcinoma). Notably, these cell lines were neither included in the STL CS nor the broader LINCS L1000 dataset, providing a stringent test of cross-context generalizability.

The STL CS demonstrates robust cross-context predictive power, showing correlation with STL001 treatment responses in both FLO1 (Spearman *ρ* = 0.728) and OVCAR8 (*ρ* = 0.574) cell lines (Fig. 3A-B). When evaluated as a binary classifier, the CS achieves exceptional performance, with AUROC values of 0.949 (FLO1) and 0.928 (OVCAR8) for predicting drug response (Fig. 3C-D). These results validate that our multi-cell-line consensus approach captures fundamental pharmacological properties that generalize to novel cellular environments beyond the training dataset.

**Fig. 3.**
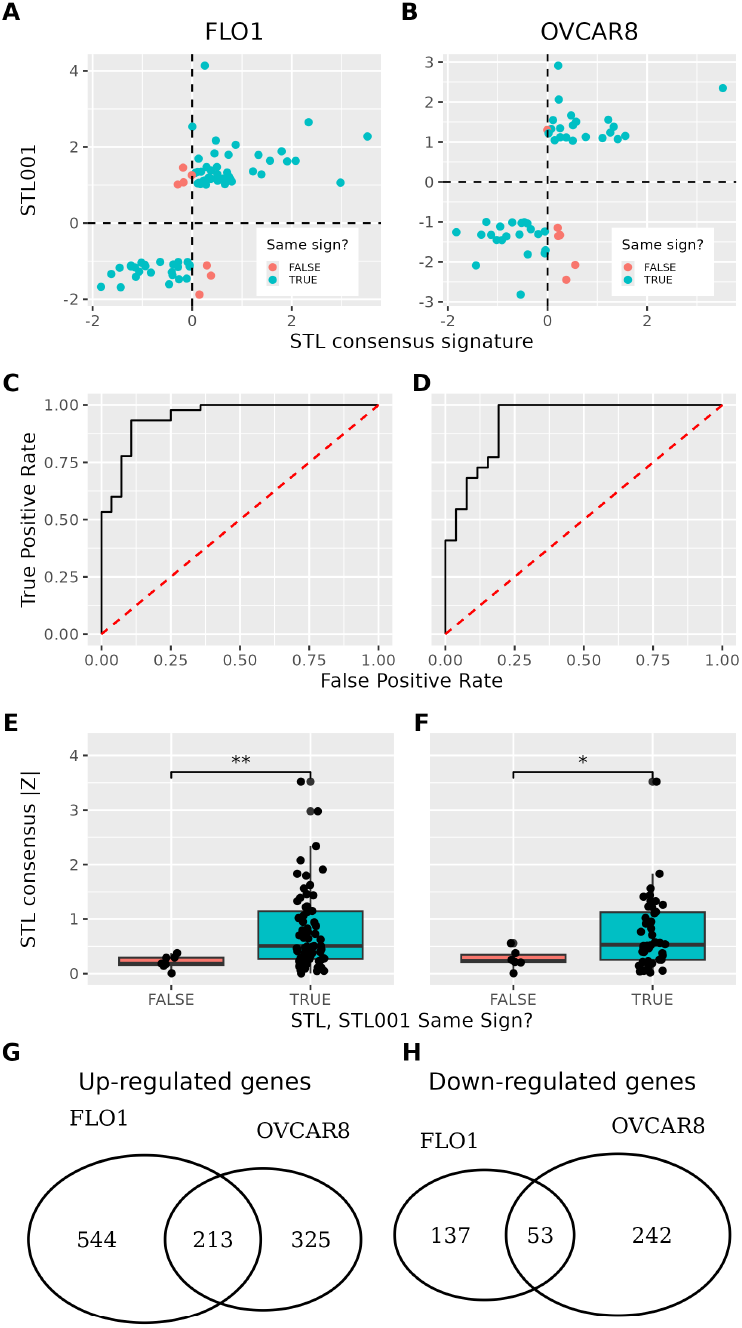
STL CS predicts responses in unseen cellular contexts. A) STL CS vs STL001 FLO1 log_2_ fold change for differentially expressed LM genes in STL001. B) STL CS vs STL001 OVCAR8 log_2_ fold change for differentially expressed LM genes in STL001. AUROC of STL001 LM gene regulation in (C) FLO1 and (D) OVCAR8 as predicted by STL CS. E) Absolute value of STL CS split by correctly and incorrectly predicted STL001 LM response in (E) FLO1 and (F) OVCAR8. Overlap of STL001induced (G) up- and (H) down-regulated genes in FLO1 and OVCAR8 cell lines.

Since our CS are expressed as Z-scores, they inherently quantify confidence in gene regulation by each perturbagen across profiled contexts. Notably, Fig. 3A-B demonstrate that most incorrect predictions occur in lower-confidence (smaller absolute Z-score) ranges, suggesting that Z-score magnitude may reflect predictive reliability in novel cellular environments. We performed one-sided Mann-Whitney U tests comparing Z-score magnitudes between correctly and incorrectly predicted genes, finding significant separation in both FLO1 (*p* = 7.24*×*10^*−*3^) and OVCAR8 (*p* = 2.48*×*10^*−*2^) (Fig. 3E-F). This analysis confirms that Z-score amplitudes are indeed predictive of cross-context regulatory accuracy.

### Imputed consensus signature genes remain predictive of new cellular contexts

Although our analyses primarily focus on LM genes, CS can be extended to encompass the full set of 12,328 genes from LINCS. To systematically evaluate predictive performance, we also compared STL CS against STL001 responses using: (1) LINCS’ best inferred gene (BING) set, and (2) lower-confidence imputed genes. As expected, the least reliable imputed genes showed poor correlation and classification performance (Table 1). In contrast, BING genes maintained strong predictive power (AUROC = 0.904 [FLO1], 0.794 [OVCAR8]), with Z-score magnitudes significantly associated with prediction confidence (*p* = 3.70*×*10^*−*20^, 4.91*×*10^*−*14^). While LINCS imputes gene expression, which is context-dependent, by averaging expression values over samples from 30 tissue types (15), CS mitigate such cell-type-specific imputation biases by averaging over several cell lines. Our findings strongly suggest that BING genes regain substantial cross-context predictive value when compiled into CS.

**Table 1.** STL CS is predictive of STL001 cellular responses across different gene sets. Comparison of STL CS to STL001 differentially expressed genes in FLO1 and OVCAR8 by Spearman correlation, AUROC, and one-sided MannWhitney U test p value. AUROC computed with STL001 response as labels and STL consensus signature as predictions. Mann-Whitney U test computed comparing consensus signature magnitude, segmented by agreement to STL001 re-sponse, null hypothesis |*Z*|_correct_ *≤* |*Z*|_incorrect_. Significant p values are in bold.

| FLO1 |  |  |  |
| --- | --- | --- | --- |
| Gene Set (Size) | $\rho$ | AUROC | PMW |
| LM (73) | 0.728 | 0.949 | **$7.24 \times 10^{-3}$** |
| BING (486) | 0.397 | 0.904 | **$3.70 \times 10^{-20}$** |
| Non-BING (61) | 0.040 | 0.718 | **$1.21 \times 10^{-2}$** |

**Table 1. STL CS is predictive of STL001 cellular responses across different gene sets.**
| OVCAR8 |  |  |  |
| --- | --- | --- | --- |
| Gene Set (Size) | $\rho$ | AUROC | PMW |
| LM (48) | 0.574 | 0.928 | **$2.48 \times 10^{-2}$** |
| BING (443) | 0.377 | 0.794 | **$4.91 \times 10^{-14}$** |
| Non-BING (58) | 0.040 | 0.593 | $1.88 \times 10^{-1}$ |

### STL001 responses in FLO1 and OVCAR8 present independent validations of STL consensus signatures

Importantly, the predictive accuracy observed in these two experimental validations is independent. While STL001 upregulates 757 genes in FLO1 and 538 in OVCAR8, only 213 are shared (Jaccard index = 0.197, Fig. 3G). Similarly, downregulated genes show minimal overlap (190 FLO1, 295 OVCAR8, 53 shared; Jaccard = 0.123, Fig. 3H). This pattern persists across gene subsets: LM genes (Jaccard = 0.196 up, 0.227 down), BING genes (0.196, 0.126), and other imputed genes (0.139, 0.074). The low overlap confirms that FLO1 and OVCAR8 provide truly independent validations, robustly demonstrating that the STL CS captures fundamental drug effects that transcend cellular context-specific responses.

Collectively, this is a proof-of-concept that CS can predict the response to perturbagens in previously untested cell lines, alleviating the potential need for further experimentation. We note that as only the differentially expressed genes were made available for these experiments that the performance we achieve may be overstated. To this end, we perform a more exhaustive evaluation of the predictive capabilities of our CS (see Supplemental Note). To our knowledge, no other hypothesis-free method is capable of predicting this broad level of gene responses to perturbations, let alone to proactively identify high- and low-confidence predictions.

### How cell-cycle genes evade shRNA knockdown

To apply our CS, we investigated the KD of LM genes and found that while most genes are successfully knocked down, three key cell-cycle genes – CCNA1, ORC1, and SOX2 – are significantly up-regulated (Fig. 4A). EnrichR analysis of these genes against all LM genes revealed significant enrichment for cell-cycle processes (Table S2), (27–29). To further validate this unexpected KD resistance phenotype, we examined the constituent LINCS CGS signatures comprising our consensus. Fig. 4B confirms that every CGS signature for these KDs has positive Z-scores across core cell lines with minimal overlap. We emphasize that our CS are derived only from on-target shRNA profiles meeting LINCS’ stringent gold-standard reproducibility criteria. Further, all considered shRNA sequences can successfully target each known wildtype isoform of the target gene, so alternative splicing is unlikely to be a confounder of our results. The immediate implication of this finding is that successful KD of these genes fails to down-regulate their expression, perhaps due to their established roles in modulating the fundamental cellular processes of cell-cycle progression and stemness maintenance.

**Fig. 4.**
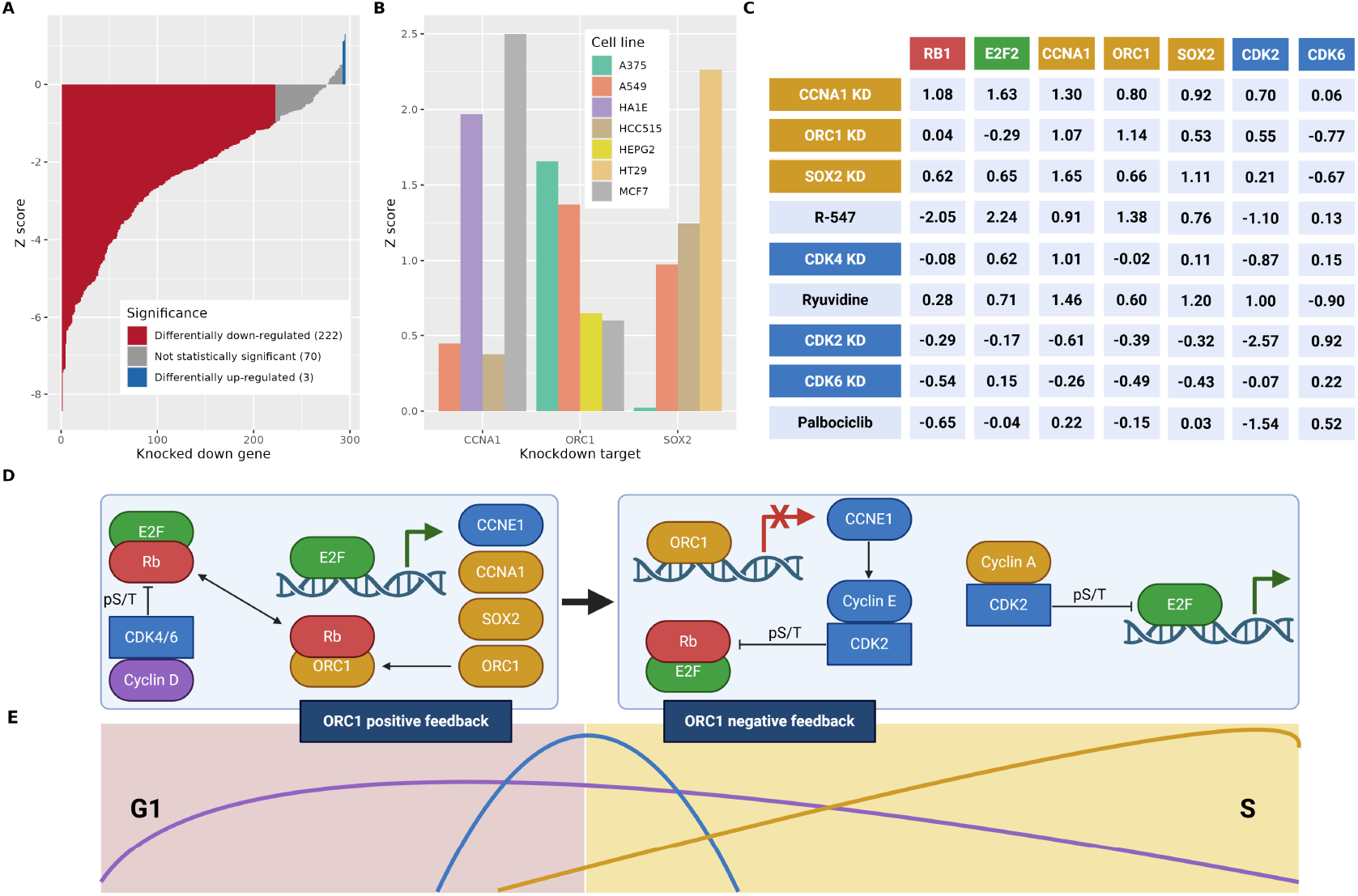
Cell-cycle resists arrests via positive and negative ORC1-mediated feedback loops to E2F. A) CS of LM genes with gold-standard KD CS, labeled by significance. B) CGS Z scores as reported by LINCS of CCNA1, ORC1, and SOX2 after their own KD in core cell lines. C) CS response to selected cell-cycle KDs and CDK inhibitors. D) Involvement of ORC1 in G1/S phase transition. CDK4/6-Cyclin D hyper-phosphorylates Rb, inactivating it and allowing the activation of E2F beginning the G1/S transition. E2F transcribes Cyclin A (CCNA1), ORC1, and SOX2 as well as CDK2, Cyclin E (CCNE1) and others. ORC1 can compete with E2F1/2 for Rb binding. ORC1 suppresses CCNE1 transcription and subsequent CDK2-Cyclin E phosphorylation of Rb. CDK2-Cyclin A phosphorylates E2F nearing end of S phase. E) Sketch of cyclin levels (D, E, A) throughout G1 and S phases. Triggered by Cyclin D-CDK4/6, ORC1 positive feedback loop increases Cyclin E and A; Cyclin E is down-regulated by ORC1 negative feedback loop during S phase. C-E created with Biorender.com.

### ORC1-mediated E2F bistability regulates G1/S checkpoint

Notably, these knockdown-resistant genes form a self-reinforcing regulatory circuit, exhibiting reciprocal upregulation (Fig. 4C). Transcriptional analysis reveals they are regulated by E2F as core components of a conserved cell-cycle signaling pathway (30–33). In G1 phase, the retinoblastoma protein (Rb) binds and inhibits E2F; CDK4/6-cyclin D-mediated Rb hyperphosphorylation then releases E2F to activate cell-cycle progression genes (34). The observed up-regulation of E2F2 following CCNA1 and SOX2 knockdown (Fig. 4C) led us to hypothesize that E2F transcriptional activity mediates the resistance phenotype. As shown in Fig. 4D, this mechanistic link is further supported by the fact that ORC1 competes with E2F for binding to Rb at overlapping sites (35, 36). This competitive binding suggests that ORC1-Rb interaction liberates E2F proteins, enabling their transcriptional activation of CCNE1, CCNA1, SOX2, ORC1 among others-a mechanism we term the ORC1 positive feedback loop. As shown by the CS of ORC1 KD and related genes (Fig. 4C), disrupting this loop by ORC1 down-regulation leads to the down-regulation of E2F2 and potentially cell-cycle arrest. Importantly, CDK2-cyclin E (CCNE1) complex also drives Rb hyperphosphorylation, committing cells to cycle progression (37). However, ORC1 and other proteins inhibit CCNE1 transcription (38), defining an ORC1 negative feedback loop (Fig. 4D-E). Thus, ORC1 knockdown emerges as the most effective perturbation to E2F activity, operating through dual mechanisms: (1) suppression of E2F2 activity (*Z* = −0.29) by blocking the positive feedback loop and (2) inhibition of CCNE1 (imputed *Z* = −0.06) expression, thereby utilizing the ORC1 negative feedback loop to further inhibit cell-cycle progression (Fig. 4D-E).

### CDK inhibitors recapitulate E2F-mediated resistance phenotype

A comprehensive analysis of CS from 12 CDK inhibitors (targeting seven distinct CDKs) reveals persistent up-regulation of CCNA1 and SOX2 (Fig. 4C, Table S3). Strikingly, the paradoxical E2F2 up-regulation by R-547 — a potent CDK2 inhibitor expected to block the negative feedback loop in Fig. 4D — validates the dominant role of the ORC1-mediated positive regulation in maintaining E2F2 activity. Notably, R-547 was discontinued following Phase 1 clinical trials.

Among the 12 CDK inhibitors profiled, palbociclib, a selective inhibitor of both CDK4 and uniquely of CDK6, stands as the sole compound exhibiting only marginal up-regulation of both CCNA1 (*Z* = 0.22) and SOX2 (*Z* = 0.03), and marginal down-regulation of both ORC1 (*Z* = −0.15) and E2F2 (*Z* = −0.04). Our CS provide compelling evidence that pharmacological inhibition of CDK4/6 blocks both E2F2 activation (therefore, ORC1 expression) and subsequent engagement of the ORC1-mediated regulatory pathways. Mechanistically, while CDK6 and CDK2 KDs produce remarkably similar transcriptomic profiles, their divergent E2F2 responses can be rationalized by their distinct compensatory up-regulation: CDK2 KD significantly increases CDK6 expression (*Z* = 0.92), whereas CDK6 KD fails to induce CDK2 (*Z* = 0.22). This asymmetric regulatory relationship suggests CDK6 occupies a privileged position in E2F2 modulation, even when compared with CDK4 inhibitors such as ryuvidine (39, 40). These findings explains the success of CDK4/6 inhibitors palbociclib, ribociclib, and abemaciclib as cancer therapies (41). Interestingly, ebvaciclib, a CDK2/4/6 inhibitor (42) in Phase 2 trials, could synergistically provide a more significant down-regulation of CDK6, with a potential better profile by blocking both the positive and negative ORC1-mediated feedback loops that regulate E2F to maintain cell cycle pro-gression.

## Discussion

This work establishes a unified framework integrating systematic biological inference with perturbation biology to uncover evolutionarily conserved mechanisms that regulate fundamental biological processes. Leveraging the LINCS L1000 database, we demonstrate that although baseline gene expression varies substantially across cellular contexts, the underlying regulatory architecture exhibits remarkable consistency with stability approaching that of technical replicates. This robustness persists across diverse perturbations (genetic and chemical), timepoints, and doses. Methodologically, we derived high-fidelity consensus signatures (CS) for over 3,500 perturbagens (small molecules and shRNAs), integrating data across these features. By transcending cellular heterogeneity, our method amplifies biologically consistent signals, enabling discovery of regulatory principles inaccessible to context-specific analyses.

We further validate our CS framework through out-of-dataset testing, demonstrating its generalizability across novel cell lines and structurally different compounds. Specifically, we show that our CS for compound STL is highly predictive of STL001 (a structural analog with similar phenotype (21)) response in two cell lines outside the purview of LINCS L1000. Strikingly, the predictive power of LM genes extends to >10,000 imputed genes (BING; Table 1), demonstrating robust scalability. This work provides a cogent framework to benchmark RNA-seq against reference transcriptomic profiles that are obscured in single-cell-line analyses. By transcending conventional log_2_-fold-change-based analysis, this hypothesis-free, genome-wide approach quantifies transcriptional response confidence across the entire gene repertoire. This paradigm shift represents a critical advancement in interpreting large-scale perturbation effects.

We demonstrate that the systematic interrogation of LINCS perturbation data provides unique mechanistic insights. CS screening across LM genes reveals that most genes respond to shRNA knockdown through expected down-regulation of their expression. Strikingly, only three genes demonstrate statistically significant resistance to their own knockdown CCNA1, ORC1, and SOX2 (Fig. 4A) all of which are established regulators of cell-cycle progression. Building on this observation, we re-evaluated the robustness of the G1/S cell-cycle transition in response to targeted perturbations of the CDK regulatory network. Our analysis reveals a novel ORC1-mediated E2F bistability that regulates the G1/S checkpoint. This finding rationalizes current therapeutic strategies (e.g., CDK4/6 inhibition), while anticipating potential failure risks (e.g., E2F2 up-regulation). More importantly, it provides actionable insights by validating ORC1 as a promising preclinical target.

Consensus signatures offer transformative advantages by integrating multi-cell-line data to: reduce noise through suppression of cell-specific artifacts (*ε*_*c*_(*C*)) while enhancing shared pharmacological signals; uncover conserved drug-class mechanisms by averaging out cell-line-specific regu-latory variants; and, normalize dose/kinetic variability that often obscures true biological responses in single-cell analyses. These capabilities offer a robust framework to enhance reproducibility and clinical relevance in pharmacogenomics. While systems biology has yet to fully decipher the context-dependent interplay of genes, proteins, and metabolites governing cellular function as a function of gene expression, our work reveals a breakthrough: perturbation biology, quantified through Z-score analysis of differential gene expression, uncovers underlying principles not of gene expression, but of fundamental cellular regulation. The consensus signatures we derive from the LINCS initiative demonstrate that aggregated cellular responses to perturbagens - including pharmacological agents and genetic perturbations - preserve both the essential features of individual experiments and their biological relevance across diverse cell types. Collectively, our findings represent a powerful strategy to decode the context-independent principles governing core biological processes, enabling hypothesis-free interrogation of how molecular perturbations drive emergent cellular phenotypes.

## Methods

### LINCS L1000 Dataset

In this work, we used small molecule experiments from the LINCS L1000 [Phase I](https://www.ncbi.nlm.nih.gov/geo/query/acc.cgi?acc=GSE92742) and [Phase II](https://www.ncbi.nlm.nih.gov/geo/query/acc.cgi?acc=GSE70138) datasets as well as short hairpin RNA (shRNA) knockdown experiments from Phase I. This data is stratified into levels. Level 3 is expression data for 978 landmark (LM) genes that have been quantile normalized, which are further used to impute expression for over 11,000 genes. The most reliable of these imputed genes form the best-inferred gene (BING) set. Level 4 is comprised of differential Z scores between the treatment and control. Level 5 combines multiple Level 4 replicates of the same experiment into a single signature.

For shRNA gene knockdowns (KDs), LINCS provides consensus genetic signatures (CGS), sometimes known as Level 6. These signatures combine multiple Level 5 signatures targeting the same gene with separate shRNA sequences to construct consensus on-target signatures. The relationships between these data levels and our consensus signatures (described below) are visualized in Fig. 5.

**Fig. 5.**
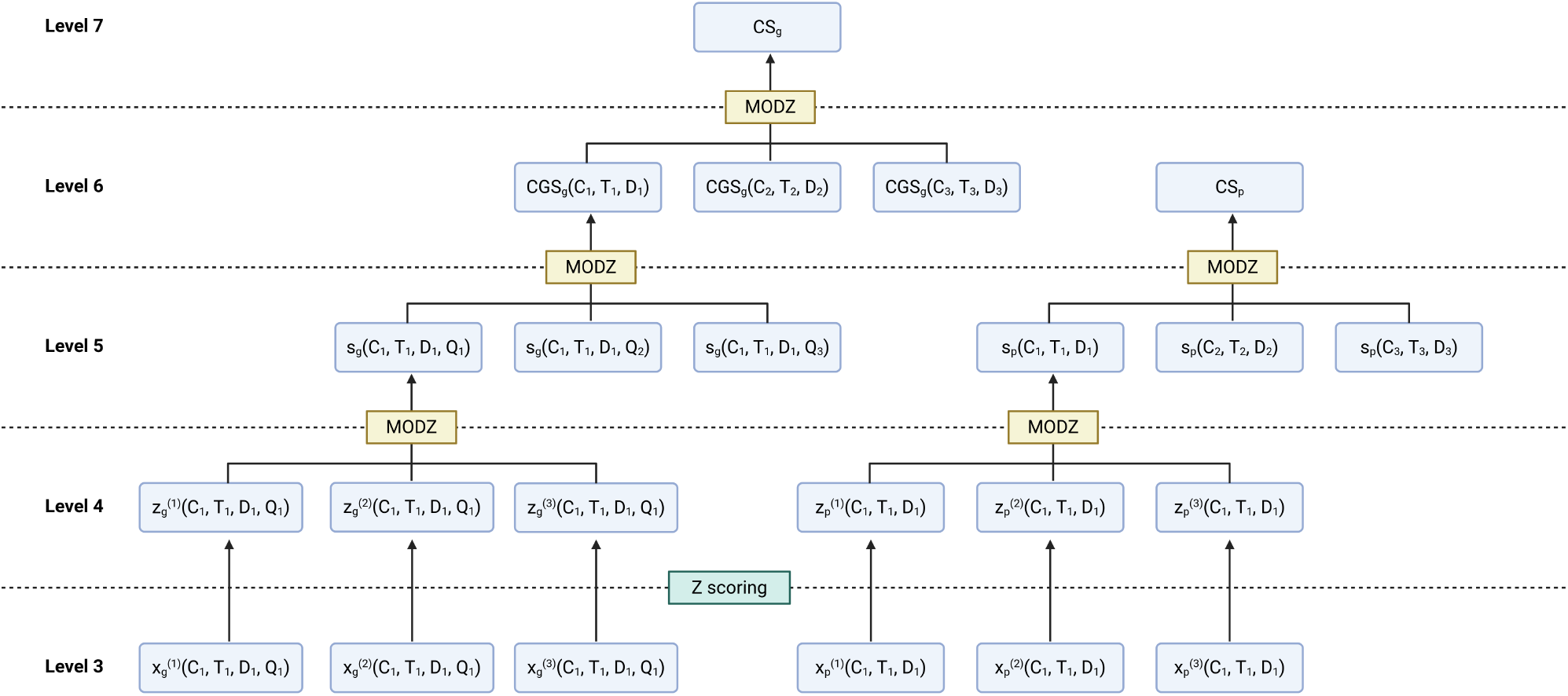
LINCS L1000 data levels and processing. Level 3 data contains replicates of gene expression after gene KD _*g*_ or small molecule *p* treatment. Level 4 data contains the same replicates but represents gene regulation by Z scores. Multiple experimental replicates are combined using the MODZ procedure to generate Level 5 signatures. Multiple KD signatures with the same cell line, dose, and time but varying shRNA sequences are combined with MODZ again to create consensus genetic signatures (CGS, Level 6). Our work contributes both consensus signatures (CS) for small molecules, compiling several Level 5 signatures with different cell lines, doses, and timepoints using MODZ, as well as KD CS that does the same but with Level 6 CGS signatures.

While most experimental assays quantify differential gene expression by log_2_ fold change, LINCS instead computes Z scores

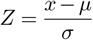

where *x* represents gene expression, and *µ* and *σ* represent the average and standard deviation of gene expression of *x*. Unlike log_2_ fold change where a larger magnitude implies a larger deviation from normal expression, in LINCS Level 4 data and up a larger Z score simply indicates a larger confidence in the regulation of the gene.

The vast majority of signatures in LINCS L1000 come from the 9 core cell lines: A375, A549, HA1E, HCC515, HEPG2, HT29, MCF7, PC3, and VCAP.

### Computation of consensus signatures

To derive a consensus perturbagen signature *CS*_*p*_ for a given perturbation *p* in the LINCS L1000 dataset, we build upon LINCS’s established consensus genetic signatures (CGS) framework (43), which suggests the following formal model:

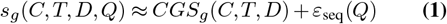

Any gene KD experiment in cell *C* with dose *D* at time *T* and shRNA sequence *Q* can be approximated by the CGS under these conditions plus *ε*_seq_(*Q*), a noise term associated with shRNA sequence to represent known off-target effects. We generalize this model to write

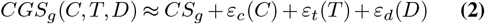

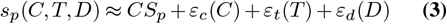

where *ε*_*c*_, *ε*_*t*_, and *ε*_*d*_ are noise terms dependent on the experimental cell *C*, timepoint *T*, and dosage *D*. To obtain CGS signatures, LINCS computed a weighted average of signatures with varying sequences *Q* as follows (43)

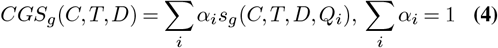

This approach generates robust on-target signatures by minimizing the sequence-specific error term *ε*_seq_(*Q*). Building on this framework, we employ LINCS’ established moderated Z-score (MODZ) weighting scheme to aggregate all available on-target signatures—whether from gene KDs (*g*) or small molecule perturbations (*p*)—into a consensus perturbagen signature (CS). The resulting CS

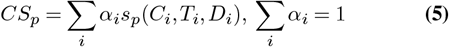

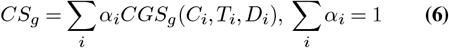

captures consistent transcriptional responses across all profiled cellular contexts, temporal conditions, and dosage regimes.

Particularly, given a set of signatures of perturbagen 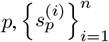, the MODZ weighting scheme is defined by

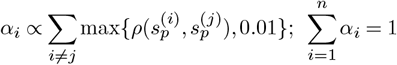

where *ρ*: ℝ^*d*^ *×* ℝ^*d*^ → [−1, 1] is the Spearman correlation coefficient, meaning experiments are weighted based on how similar they are to other experiments. Notably, the Spearman correlations are thresholded to 0.01, forcing every experiment to have a positive, nonzero weight and factor into the final signature. Further, when extending CS to entail imputed genes, we only consider the LM genes for the correlations above (*d* = 978), but apply the weights to all genes as outlined by Himmelstein et al. (44).

This procedure was previously used to combine small molecule experimental replicates (Level 4) into Level 5 signatures, as well as to create the CGS signatures from Level 5 shRNA signatures. In this work, we use the same procedure to combine Level 5 small molecule signatures into small molecule CS and Level 6 CGS signatures into KD CS. Importantly this is not a traditional meta analysis, where the absolute value of the Z score would increase given a larger burden of proof (larger *n*). We opted for MODZ over options such as Stouffer’s method so that the Z scores for perturbagens with large differences in *n* could be more readily compared.

We did not create consensus signatures for every available perturbagen. We excluded any perturbagen that did not have experiments in at least four distinct cell lines. We also mandated that the compiled signatures met the LINCS goldstandard requirement for self-consistency, specifically that the 75th percentile of pairwise Spearman correlations was at least 0.2. LINCS also usually requires meeting a distinctness test threshold to be considered gold-standard, but they omitted this requirement for the CGS signatures. Given that we are also combining signatures with varied experimental considerations, we dropped this requirement as well.

### Random signatures

To determine whether the MODZ procedure artificially homogenizes signatures regardless of biological content, we computed inter-perturbagen correlation for randomly generated signatures representing increasing numbers of “cell lines”. For each of the nine core cell lines, we summarized gene expression of common perturbagens by a normal distribution and randomly generated samples from these distributions to create “cell-line-specific” random signatures. We then combined increasing numbers of these random signatures to form random signatures that represented a specific number of cell lines and computed inter-perturbagen similarity using Spearman correlation.

### Intra-class correlation

We investigated whether drugs in the same pharmacological class were more or less similar according to consensus signatures versus experiments in single-cell lines. We limited our analysis to the 9 core LINCS cell lines. Classes are taken from the 16 most populated classes from the LINCS data portal (26). We removed quaternary ammonium derivatives from our analysis, as only a single compound was profiled in at least 4 of 9 core cell lines.

To measure intra-class similarity captured by consensus signatures, we created consensus signatures using only experiments from the 9 core cell lines and computed the pairwise Spearman correlation between each pair of perturbagens in each class. For single-cell-line intra-class similarity, for each pair of drugs in a given pharmacological class and cell line, we computed the Spearman correlation between every pair of signatures. Finally for each combination of pharmacological class and cell line, we compute a 1-sided Mann-Whitney U test comparing the single-cell and consensus signature correlations. Fig. 1H reports the mean correlation in each setting and Table S1 reports all p-values.

## Data Availability

Data available upon request.

## ACKNOWLEDGEMENTS

We want to thank Dr. Andrei Gartel and Dr. Sanjeev Raghuwanshi for providing STL001 RNA-seq data in FLO1 and OVCAR8 cell lines, and Dr. Maria Chikina and Dr. Wayne Stallaert for helpful advice. Supported by NCI 5R01CA259124-04.

