## Supplemental Note, Figures for "Cell-type-agnostic differential gene expression uncovers conserved principles of cellular regulation"

#### Prediction of new LINCS 2020 signatures

Beyond our validating that STL CS predicts STL001 response, we also performed a more systematic validation of our CS on the newest release of the LINCS L1000 dataset - LINCS 2020. As we previously demonstrated that lower-confidence CS predictions were more often incorrect ones, we only evaluated the predictions from the differentially expressed genes (DEG) of a CS. After removing experiments already found in LINCS phase I and II, we divided the new signatures into three broad categories of increasingly difficult validations. First, we considered whether a perturbagen's CS was predictive of cells, times, and dosage ranges that were already represented in the CS. Across 1,488 drugs with new such data, we found on average that our CS could predict the entire signature quite well when treated as a binary classifier as before (median AUROC = 0.803, Figure S1). When we further restricted our predictions to genes that are differentially expressed both in the CS and the new signature, we do incredibly well (median AUROC 0.931, Figure S1). Over 96% of drug CS were better than random (AUROC > 0.5) predictors for new signatures from familiar contexts, even if the specific cell line-timepoint-dosage combination had not been seen together.

Next we tested if our CS could extend to a singular new context (cell line, time point, or dose) when the other variables had been profiled before. We were able to profile 843 drugs in a new cell line at previously seen doses and times, 516 drugs at a new dose in previously seen cells and times, and 135 drugs at a new time in previously seen cells at similar doses. We observed a drop-off in predicting all CS DEG (median AUROC = 0.689, 0.694, 0.717 for new cell lines, doses, and time points, respectively, Figure S1), although DEGs conserved across contexts were still predicted well (median AUROC = 0.829, 0.837, 0.831, Figure S1). These results indicate that our CS may not be able to generalize as easily to new contexts as we would expect given our previous validation with STL001. Yet, given the performance on the intersection of DEG, we infer that DEGs generally do not flip regulation in a new context, further solidifying our general hypothesis that the gene regulatory response of a perturbagen is relatively well conserved. Additionally, we find that our CS are better than random predictors of response in a new context for 87%, 87%, and 94% of drugs profiled in a new cell line, dose, and time point respectively.

Finally, we tested how our CS could predict signatures of a drug in entirely unseen cell lines, time points, AND doses at the same time. For this test, we had 82 drugs to evaluate on. Here we observe a steep drop in performance for predicting the entire signature (median AUROC = 0.622, Figure S1) as well as the DEG (median AUROC = 0.720, Figure S1). While disappointing, this result affirms the complexity of the biological systems we are interrogating as well as highlights the shortcomings of a simplistic linear model for predicting perturbagen response in a zero-shot classification setting. Yet even in this hardest-case prediction scenario, we demonstrate better-than-random predictions for nearly 80% of drugs. Thus, our CS may represent a useful baseline for

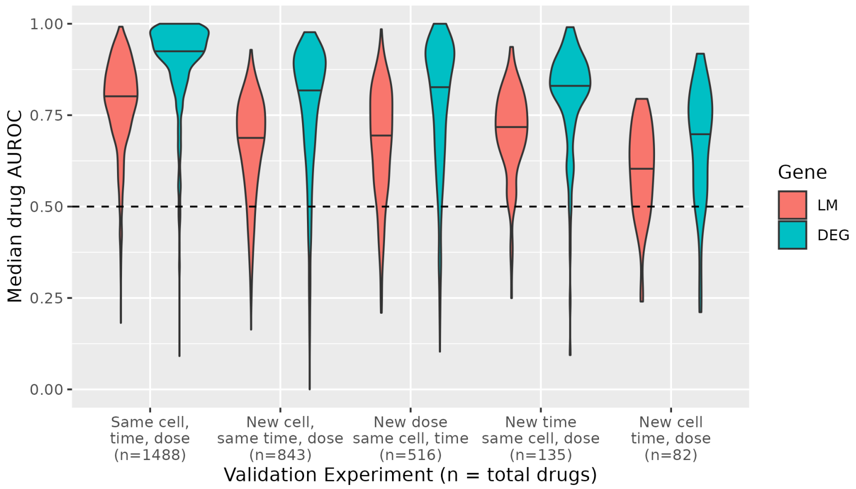


Figure S1: **Consensus signatures (CS) are predictive of held-out data from LINCS 2020 update.** Evaluation of CS as binary classifier on new data from LINCS L1000 2020 update. Only differentially expressed genes (DEG) from CS used to make predictions. Fill corresponds to prediction accuracy on all CS DEG and DEG for both CS and new signatures, respectively.

further studies attempting to predict perturbation response with more complex architectures.

We note that the LINCS 2020 release did not include any additional shRNA knockdown experiments, instead favoring CRISPR knockouts (KOs). We opted not to compare our KDs to KOs, as they are fundamentally different perturbations. KDs utilize the cell's built-in RNA interference pathways to silence expression at the level of mRNA transcripts, whereas KOs delete a gene from a cell's DNA, preventing any possible expression of that gene. This is an important difference as KDs can potentially be resisted by autoregulatory feedback mechanisms whereas KOs can not.

#### Cell-cycle phase annotation

Given that we are considering cell-cycle perturbations, it is important to consider cell-cycle phase as a factor in our results. The cell cycle is known as a profound confounder of transcriptomic response [3, 2], so we verified whether the LINCS signatures for these small molecules and KDs discussed above indicate a preference for a specific cell-cycle phase. Particularly, we looked at the expression of marker gene pairs from Scialdone et al. and found that indeed, several of the known CDK inhibitors exhibited phenotypes associated with G1 phase, most notably the FDA-approved palbociclib (Figure S2). By contrast, gene knockdowns should either no obvious phase preference or a weak preference for G1 (CDK4, SOX2, and ORC1). While this marker pairs method was demonstrated to recapitulate

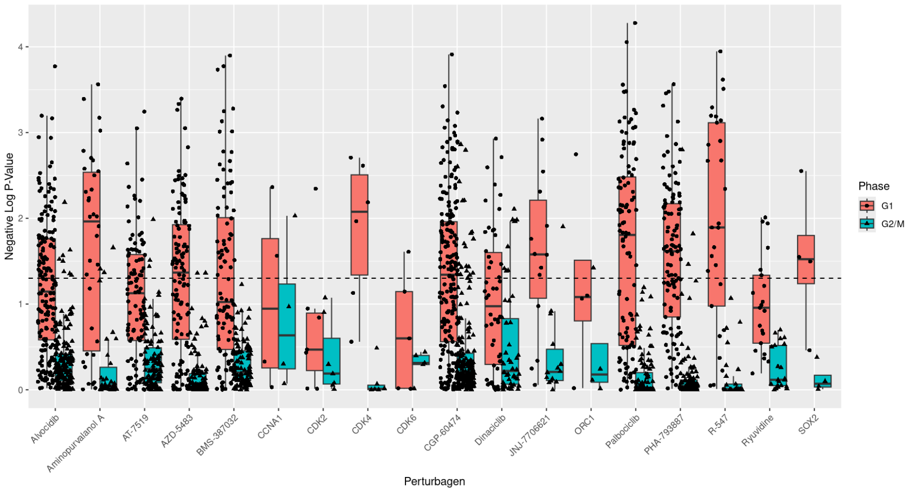


Figure S2: **Chemical perturbations, not genetic perturbations, to cell-cycle shows arrest phenotypes.** Eight marker gene pairs predictive of G1 and nine marker gene pairs predictive of G2/M were measured in LINCS signatures for 12 CDK inhibitors and 6 cell-cycle-relevant KDs. The difference of these marker genes was measured for each experiment and summarized by the negative log p-value of a one-sided, one-sample t-test. Several CDK inhibitors show evidence of strong preference for G1 (AZD-5483, BMS-387032, Palbociclib, PHA-793887, R-547), whereas knockdowns do not show as strong preference for G1. Marker gene pairs taken from [1], see Table S1.

the phases of staged cells measured with bulk RNA-seq [1], our results are far less conclusive; 1) our only reference was the reported top 10 disjoint gene pairs for each phase, 2) we could only actually utilize 8 pairs for G1 and 9 for G2/M, with only 3 pairs in total being both LM genes, so samples are small and include potential imputation biases (see Table S1), and 3) it is unclear if the LINCS CGS signatures entangles any latent cell-cycle phenotypes by averaging over different shRNA signatures. However these results are an indication that while the phenotypes embodied by the CDK inhibitors may be linked to cell-cycle arrest in G1, this may not be the case for the knockdowns we considered. This further contextualizes our proposed mechanisms of cell-cycle resilience as being independent of whether cells are arresting or not.

| G1 marker gene pairs | | | |
|---|---|---|---|
| Gene 1 | LINCS space | Gene 2 | LINCS space |
| HMGCR | LM | CCNB1 | LM |
| B4GALT1 | BING | PLK1 | LM |
| ERH | BING | UBE2C | LM |
| CSNK2A2 | LM | CDC25C | BING |
| FOS | LM | FZR1 | BING |
| WTAP | BING | TACC3 | BING |
| SEC62 | BING | KIF23 | BING |
| ARGLU1 | BING | TUBB4B | BING |
| **G2/M marker gene pairs** | | | |
| Gene 1 | LINCS space | Gene 2 | LINCS space |
| CCNB1 | LM | USP22 | LM |
| NUSAP1 | LM | PCNA | LM |
| UBE2C | LM | PCNP | BING |
| PLK1 | LM | DCTPP1 | BING |
| CDK1 | LM | SON | BING |
| PRC1 | BING | WAPL | BING |
| ARL6IP1 | BING | TIPIN | BING |
| BUB3 | BING | LARP4 | BING |
| TPX2 | BING | PPP1CB | BING |

Table S1: **Marker gene pairs for G1, G2/M cell cycle phase annotation**. Pairs taken from top 10 most important, disjoint features of marker gene pairs model from [1], ordered by gene set, feature importance. For given pair $(g_1, g_2)$ and phase, was found that $g_1 - g_2 > 0$ was significantly more likely in given phase than other and thus predictive of phase. Two G1 and one G2/M marker pair is not included as one gene in pair not included in LINCS L1000 data. Landmark (LM) genes are measured in every experiment, and are the most reliable markers. Best-inferred gene (BING) set are imputed from LM gene expression and subsequently less reliable markers but included to indicate larger trends.

### Supplemental Figures

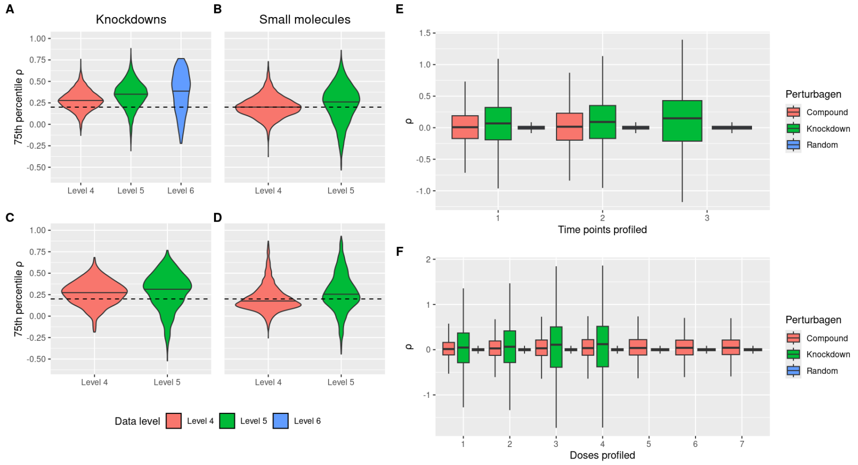


Figure S3: **Consensus perturbagen signatures reveal consistency across time scales, doses.** A) Small molecule intra-dose (replicate) Z score correlation and inter-dose signature correlation for small molecules profiled at at least 2 doses. B) shRNA KD intra-dose (replicate) Z score correlation and inter-dose CGS signature correlation for KDs profiled at at least 2 doses. C) Small molecule intra-time (replicate) Z score correlation and inter-time signature correlation for small molecules profiled at at least 2 time points. D) shRNA KD intra-time (replicate) Z score correlation and inter-time CGS signature correlation for KDs profiled at at least 2 time points. E) Inter-perturbagen correlation for CS and random signatures utilizing varying numbers of doses. Small molecule doses 10.00 $\mu$M, 5.00 $\mu$M, 3.33 $\mu$M, 1.11 $\mu$M, 0.37$\mu$M, 0.12 $\mu$M, 0.04 $\mu$M. Knockdown doses 5.0 $\mu$L, 2.0 $\mu$L, 1.5 $\mu$L, 1.0 $\mu$L. F) Inter-perturbagen correlation for CS and random signatures utilizing varying numbers of time scales. Small molecule time scales 6 h, 24 h. Knockdown time scales 96 h, 120 h, 144 h.
